# The fire blight pathogen *Erwinia amylovora* enters apple leaves through naturally-occurring wounds from the abscission of trichomes

**DOI:** 10.1101/2024.10.10.617712

**Authors:** Felicia Millett, James Standish, Jules Scanley, Katelyn Miller, John Inguagiato, Nubia Zuverza-Mena, Maritza Abril, Victoria Robinson, Yan Li, George W. Sundin, Quan Zeng

## Abstract

- The plant epidermis is a single layer of cells covering all plant organs. How pathogens overcome this barrier and enter plants is an important aspect of plant-pathogen interactions. For bacterial plant pathogens, known entry points include natural openings such as stomata, hydathodes, and mechanical injuries caused by insect feeding, wind damage or hailstorms.
- Here, we report that the fire blight pathogen *Erwinia amylovora* enters apple leaves through naturally-occurring wounds caused by the abscission of trichomes during the course of leaf development.
- Through macroscopic and microscopic observations, we depicted a clear invasion path for *E. amylovora* cells, from epiphytic growth on glandular trichomes (GT) and non-glandular trichomes (NT), to entry through wounds caused by abscised trichomes, into the epithem, and subsequent spread through xylem. We further observed that GT and NT undergo an abscission process, and that the amount of naturally-occurring wounds during abscission is associated with the increase in *E. amylovora* population. Key genes important for the colonization of GT and NT were identified. Contribution of the type III secretion system and amylovoran biosynthesis during GT colonization was validated.
- Our findings propose a novel host entry mechanism of plant pathogenic bacteria through naturally-occurring wounds during abscission of plant surface structures.

## INTRODUCTION

While some plant pathogens reside strictly on plant surfaces, most spend a significant part of their life cycle internally within plants to cause disease infections. These internal colonization sites include the apoplast, intracellular spaces, and plant vascular system elements such as xylem and phloem. Plants are covered by an epidermis layer composed of cutin, wax, and polysaccharides. The epidermis not only prevents water loss and controls gas exchange, but also acts as the first layer of innate immunity to prevent pathogen entry. Therefore, how plant pathogens overcome this protective barrier to enter hosts is a critical aspect of plant-pathogen interactions and subsequent disease infection.

Unlike fungal pathogens, bacterial plant pathogens do not encode penetration structures and therefore rely on either natural openings or mechanical injuries for their entry. Known entry points of bacterial plant pathogens include stomata (Zeng *et al*., 2010), hydathodes (Cerutti *et al*., 2017), root hairs (Perrine-Walker *et al*., 2007), lenticels (Bartz *et al*., 2016), weather-induced mechanical injuries (Dow *et al*., 2017; Pedroncelli & Puopolo, 2023) and injuries caused by insect feeding(Grafton-Cardwell *et al*., 2013). Bacterial plant pathogens have evolved mechanisms to identify the plant locations to facilitate their entry. For example, the bacterial wilt pathogen *Ralstonia solanacearum* utilizes chemotaxis to recognize exudates released through natural openings and wounds on roots (Tans-Kersten *et al*., 2001; Yao & Allen, 2006), and the crown gall pathogen *Agrobacterium tumefaciens* is chemotactic towards the plant wound chemical acetosyringone to identify and move to entry points (Stachel *et al*., 1985). *Pseudomonas syringae* produces the phytotoxin coronatine to inhibit a jasmonic acid-induced stomata closure of plants, therefore enabling successful entry (Zheng *et al*., 2012).

Prior to host entry, an epiphytic growth phase is often important for many bacterial plant pathogens. The epiphytic colonization often occurs at structures that produce secretions, such as hydathodes, stomata, and trichomes (Schlechter *et al*., 2019). Secretions produced from these structures provide not only nutrients to support pathogen growth, but also chemical signals for host recognition (Yao & Allen, 2006), and induction of bacterial virulence (Cui *et al*., 2021).

Among different plant surface structures, trichomes, the outgrowths of plant epidermis, play diverse and important roles in plant physiology, defense, and interactions with the environment (Huchelmann *et al*., 2017; Karabourniotis *et al*., 2020; Schuurink & Tissier, 2020). Based on the secretory functions, trichomes are categorized into glandular trichomes (GT) and non-glandular trichomes (NT). GTs are multicellular structures composed of differentiated basal, stalk, and apical cells that secrete substances such as essential oils, resins, and other secondary metabolites (Schuurink & Tissier, 2020). NTs are single- or multi-cellular structures with a pointed or hair-like appearance without secretion functions (Karabourniotis *et al*., 2020). On apple leaves, GTs are present on leaf serrations and along the veins, while NTs (leaf hair) are present in heavy abundance on both abaxial and adaxial sides of leaves.

Fire blight is a devastating disease of plants belonging to the Rosaceae family, such as apple (*Malus* x *domestica*) and pear (*Pyrus* species) (van der Zwet *et al*., 2012; Pedroncelli & Puopolo, 2023). The causal agent, *Erwinia amylovora*, is an enterobacterium that is listed as one of the ten most important plant pathogenic bacteria (Mansfield *et al*., 2012). In an orchard, fire blight occurs in two stages. First, the blossom blight stage occurs in early spring, during which the overwinter inoculum in the form of ooze produced from cankers is transferred to flowers, leading to necrotic, wilt and blight symptoms of flowers (Slack *et al*., 2022). Second, the shoot blight stage occurs from mid-spring to mid-summer, during which *E. amylovora* cells in ooze droplets produced from the infected flowers are spread to the newly expanded succulent shoots and causes infections (Pedroncelli & Puopolo, 2023). Infected shoots can produce more ooze, which may result in disease epidemics when environmental factors favor infection (Slack *et al*., 2017).

Like many bacterial plant pathogens, *E. amylovora* lacks cuticle penetration ability, and therefore has to enter hosts through natural openings or wounds. During the flower infection phase resulting in blossom blight symptoms, it is known that *E. amylovora* enters host flowers through nectarthodes, the natural openings at the hypanthium that secrete nectar (van der Zwet *et al*., 2012). It also has been shown that prior to host entry, *E. amylovora* grows epiphytically on the flower stigma surface (Kharadi *et al*., 2021; Zeng *et al*., 2021; Slack *et al*., 2022). This epiphytic growth stage is essential for the subsequent invasion as it establishes a large pathogen population and primes pathogen virulence (Cui *et al*., 2021; Zeng *et al*., 2021).

Compared to blossom blight, how *E. amylovora* overcomes the epidermis and enters the host during shoot blight infection is not clearly understood. According to the American Phytopathological Society Plant Disease Profile of Fire Blight, *E. amylovora* from bacterial ooze enter shoots through wounds created by feeding of piercing-sucking insects, such as plant bugs and psylla, or through injuries caused by severe weather events such as gusty wind, rain or hailstorms (Johnson, 2015). However, growers often observed shoot blight infections independent of these severe weather events or noticeable insect activities. Early research conducted to determine whether fire blight infection can occur on apple leaves without artificial injuries resulted in inconclusive results (O’Gara, 1912; Hotson, 1916; Brooks, 1926; Miller, 1929; Rosen, 1929; Tullis, 1929; Pierstorff, 1931; Crosse *et al*., 1972). Currently, most researchers use a scissors-cutting method to inoculate *E. amylovora* into shoots for virulence testing (van der Zwet *et al*., 2012).

Unlike flowers that produce various nutrient rich exudates, leaves of rosaceous plants are generally nutrient poor and do not support heavy epiphytic populations of *E. amylovora*. Norelli and Brandl reported that *E. amylovora* could colonize hydathodes and glandular trichomes of young leaves (Norelli & Brandl, 2006). Bogs et al (1998) and Campbell et al (2001) reported that *E. amylovora* is able to colonize leaf hairs (non-glandular trichomes) (Bogs *et al*., 1998; Campbell *et al*., 2001). Tullis proposed that stomata on apple leaves are entry points for *E. amylovora* (Tullis, 1929). Not only do some of these findings contradict to each other, there is also a lack of experimental evidence as to whether the epiphytic colonization of any of the above structure(s) actually leads to leaf infection.

In this study, we used a GUS-labeled *E. amylovora* strain to determine the epiphytic colonization sites of *E. amylovora* on apple leaves and to track the invasion path during shoot blight infection. We found that *E. amylovora* colonizes GTs and NTs on apple leaf surfaces. We further provided evidence that the GTs and NTs naturally abscise during leaf development, which provides opportunities for *E. amylovora* to enter the host. Finally, a transcriptomic analysis identified differentially-expressed genes in *E. amylovora* on GTs and NTs. Through mutation and GUS labeling, we confirmed that the type III secretion system and amylovoran biosynthesis are important for the GT colonization and infection by *E. amylovora*.

## MATERIALS AND METHODS

### Apple shoot inoculation and virulence assay

Two-year old bare root apple trees ‘Simmons Gala’ on G41 rootstock were purchased from Adams County Nursery and were potted in 5 gallon pots containing a mixture of 2:1:1 (vol/vol) of Promix BX (Quebec, Canada), topsoil and sand. Trees were defoliated and were maintained in an environmentally controlled growth chamber (Percival PGC 105; Perry, Iowa) prior to the experiment. *E. amylovora* strain *Ea*110 (Zhao *et al*., 2005) was cultured in Lysogeny broth (LB) overnight at 28°C. Bacteria cells were pelleted by centrifugation at 5,000 rpm for 10 min. Concentrations of bacterial suspension were adjusted to 1 x 10^8^ colony-forming units (CFU) / ml in distilled water (optical density at 600nm = 0.1). Newly expanded shoot tips approximately one to three weeks old were dipped in bacterial suspension for 30 seconds. Inoculated trees were kept at 25°C and 99% relative humidity, 14-hour days with 350 umol light. Symptoms were monitored daily.

### Detecting of *E. amylovora* presence using GUS staining

*Ea*1189*::gus*, an *E. amylovora* strain with *gus* gene integrated into the chromosome (Mukhtar *et al*., 2024)was dip-inoculated to intact apple leaves using the same method described above. Inoculated leaves were collected at 12, 24, 48, and 36 hrs post inoculation for a GUS staining process. Leaves were submersed in acetone on ice for 15 min, then vacuum infiltrated for three 10 min intervals with 40 mg/μl X-Gluc substrate (5-bromo-4-chloro-3-indolyl-beta-D-glucuronic acid, cyclohexylammonium salt in dimethylformamide), 0.1 M potassium ferricyanide K_3_[Fe(CN)_6_], 0.1 M potassium ferrocyanide K_4_[Fe(CN)_6_], 1 M sodium phosphate NaPO_4_, 0.5M ethylenediaminetetraacetic acid (EDTA) and distilled water) and incubated at 37°C overnight in the dark. Twenty-four hours later, leaves were placed in 80% ethanol for 40 min at 60°C, followed by 40 min in 95% ethanol at 60°C for de-staining of the chlorophyll. They were then transferred to 50% ethanol at room temperature and observed with a Zeiss Discovery V12 Stereo microscope (Carl Zeiss Inc., Oberkochen, Germany). Images were acquired through with a digital AxioCam HRc camera and AxioVision software, version 4.8.1.

### Transmission electron microscopy

The method of preparing leaf tissue for transmission electron microscopy by (Cerutti, 2017) has been modified for our purposes. Leaf tissue samples (3-4 mm^2^) were cut with a razor and vacuum fixed for 10 min in 2.5% glutaraldehyde in 0.1 M sodium cacodylate buffer (pH 7.2) and 0.1% Triton X-100. Samples were then fixed for 1 hr in 2.5% glutaraldehyde in 0.1 M sodium cacodylate buffer (pH 7.2), followed by a sodium cacodylate rinse. Samples were postfixed in 2% osmium tetroxide in 0.2 M sodium cacodylate buffer for 1 hr, followed by another sodium cacodylate rinse. Samples were dehydrated with an ethanol series (25%, 50%, 70%, 95%, and 100%) and rinsed in absolute acetone three times. Samples were infiltrated in 1:3, 1:1 and 3:1 ratios of Embed 812 resin to acetone and three changes of 100% EMbed812 resin before final polymerization at 60°C for 48 hrs.

Semithin sections (1 µm) were cut with a Histo 45° Diatome™ diamond knife on a Leica Ultracut UCT microtome, mounted on glass slides and stained in a 1% toluidine blue at 60°C for 1 min. Sections were examined at the light microscope level to identify suitable material for electron microscopy. For transmission electron microscopy, ultrathin (100 nm) sections were cut with an ultra 45° Diatome™ diamond knife on a Leica Ultracut UCT microtome and collected on formvar support film slots 2×1 mm cu grids (EMS). Sections were counterstained with 2% ethanolic uranyl acetate, rinsed with distilled water, stained with 2.5% Sato’s lead citrate, and rinsed again with distilled water. Images were acquired using a Nikon alphaphot YS microscope (Nikon Instruments, Tokyo, Japan) for thin sections and a Hitachi HT7800 transmission electron microscope (Hitachi Ltd, Tokyo, Japan) at 80kV for ultrathin sections.

### Scanning electron microscopy

Apple leaf tissues (cultivar ‘Gala’) inoculated with *E. amylovora* strain *Ea*110 were collected at 3 dpi and fixed in paraformaldehyde/ glutaraldehyde (2.5% of each compound in 0.1 M sodium cacodylate buffer) (Electron Microscopy Sciences, Hatfield, PA) at 25 C overnight. Fixed tissues were dehydrated in 25, 50, 75, and 90% ethanol for 1 hr each and in 100% ethanol three times for 30 min each. Dehydrated samples were air-dried at room temperature and mounted on aluminum mounting stubs. Images were captured using a Zeiss Sigma VP FESEM (Carl Zeiss Inc., Oberkochen, Germany).

### Confocal microscopy

*Ea*1189::*gfp*, an *E. amylovora* strain with the *gfp* gene integrated into its chromosome, was constructed using a previously reported λ red recombinase method to replace the intergenic region (2425167-2426058) with a *gfp* gene. The intergenic region is located between a plasmid-like protein gene (EAM_2245) and a hypothetical protein gene (EAM_2246) of the *E. amylovora* Ea1189 genome. *Ea*1189::*gfp* was inoculated into apple leaves (cultivar ‘Gala’) using the same dip-inoculation method above. Samples were collected at 2 dpi. Cut pieces of leaves were mounted to glass slides and observed with a Leica TCS SP5 confocal microscope (Leica Microsystems, Wetzlar, Germany) with a 488-nm argon laser set for excitation of GFP while red autofluorescence spectra was recorded at 561-nm excitation for the detection of plant tissue. Image acquisition was captured with Leica LAS AF software.

### Determination of *E. amylovora* population on GT over three timepoints

Shoots of ‘Simmons Gala’ trees were dip inoculated in a suspension of rifampicin resistant *E. amylovora Ea*110 cells at a concentration of 1 x 10^7^ CFU/ml, and were maintained in at 25°C, 99% relative humidity and 14-hr days in a plant growth chamber. Samples were collected at 10 hpi, 48 hpi and 72 hpi. For each time point, 12 GTs were collected from each of 4 leaves, for a total of 48 GTs for each timepoint. GTs were collected using a disposable pipette tip and homogenized in 50 µL sterile distilled water in a microcentrifuge tube. The *E. amylovora* population was determined by plating on LB agar plates supplemented with 50 μg/ml cycloheximide and 10 μg/ml rifampicin and counting the CFU. Means separation lettering based on multiple comparison tests on pairwise rankings based on a nonparametric Kruskal-Wallis test (*P* =0.0041 for Chi-square null hypothesis of equal rank sums). Statistical analysis performed in SAS using proc npar1way wilcoxon DSCF. Box plot created in RStudio using ggplot. Means separation lettering based on multiple comparison tests on pairwise rankings based on a nonparametric Kruskal-Wallis test (*P* =0.0041 for Chi-square null hypothesis of equal rank sums).

### Quantification of *E. amylovora* abundance on NTs using qPCR

Apple leaves ‘Gala’ were collected three days after *E. amylovora* dip-inoculation (10^7^ CFU/ml). Collected leaves were dipped in liquid nitrogen to facilitate easy removal of NT with a small scalpel and were subject to DNA isolation using Qiagen DNeasy Plant Mini Kit (Cat. No. 69104). The abundance of *E. amylovora* on trichomes collected from un-netted and netted trees was quantified by determining the cycle threshold (CT) value of the *E. amylovora*-specific gene *amsC* (Pirc *et al*., 2009). qPCR of *amsC* gene was performed using a SsoAdvanced universal SYBR Green supermix (Bio-Rad, CA, USA). The *amsC* copy number was normalized by the copy number of the apple tubulin gene (Zeng *et al*., 2023) using the ΔΔ*ct* method (Livak & Schmittgen, 2001).

### Determination of NT density on leaves

Non-glandular trichome (NT) density was quantified by acquiring leaf images with Zeiss Discovery V12 Stereo microscope with a digital AxioCam HRc camera and AxioVision software and then counting the number of individual NT intersecting a 2 mm line. Data collected from two locations on the abaxial surface and two locations on the adaxial leaf surface was averaged for 15 leaves of each cultivar. Comparisons were performed using a two-sample t-test (P < 0.0001). Statistical analysis performed and boxplot created in RStudio using ggplot2, ggpubr and rstatix.

### Grayscale values as an indication of NT density

The second and third open leaves from the tip of each shoot were collected for analysis. Leaves were observed under a Zeiss Discovery V12 Stereo microscope at 50% brightness and images acquired with digital AxioCam HRc camera and AxioVision software. Configurations in AxioVision were the same for each leaf: exposure was set to 35.0 ms, white balance color offset was se to -0.10, and within display settings, brightness value was -0.40, contrast value was 1.01, and level was 1.00. Microsoft Image Composite Editor (ICE) software was used to stitch together images and reconstruct individual leaves. ImageJ (version 1.53h) was used to find the grayscale value, within a pixel value range from “0” for black, and “255” for white after threshold color configurations were set for hue (0/255), saturation (0/255), and brightness (30/255). For each leaf, grayscale value was calculated as the mean of the abaxial and adaxial leaf surface grayscale values (see Supplementary Fig. 3).

Plant material for grayscale analysis for NT density included the heirloom cultivars listed above and additional fresh shoot cuttings, obtained from the Plant Genetic Resources Unit (PGRU) in Geneva, New York, a division of the United States Department of Agriculture, Agricultural Research Service. *Malus domestica* accessions included: ‘Circassian Apple’, ‘Spartan’, ‘Milton’, ‘Hotle Rome’, ‘Ingol’, ‘Braeburn’, ‘Rhode Island Greening’, ‘Golden Delicious’, ‘Macoun’, ‘Sunrise’, and ‘Jonathan’. Accessions of *Malus* hybrids included: ‘Nipissing’, ‘Hopa’, ‘Scugog’, PRI 1744-1, and FORM 181 (35-01). Cuttings of crabapples included *M. baccata* ‘Nertchinsk’, (Siberian crabapple) and one *M. fusca*, GMAL 2841, (Oregon crabapple). Cuttings of two accessions of the wild apple, *M. sieversii* were obtained, FORM 43 (34-01) and KAZ 96 01-01P-20. A non-parametric Kruskal-Wallis test was performed to determine significant difference between means. Statistical analysis performed in SAS using proc npar1way wilcoxon DSCF. Boxplot created in RStudio with ggplot.

### Transcriptomic analysis of *E. amylovora* on trichomes

Young leaves (1 week after they unfold) of apple cultivar ‘McIntosh’ were dip-inoculated with a suspension of E. amylovora strain Ea110 in 0.5X PBS at the concentration of 108 CFU/ml. The remaining inoculum was frozen in liquid nitrogen and used for RNA isolation as the ‘Inoculum’ sample. The inoculated trees were maintained in a greenhouse. Glandular and non-glandular trichomes were harvested four days post inoculation. Glandular trichomes were collected by dissecting from the leaf serration with scissors; the non-glandular trichomes were collected by dipping the leaf in liquid nitrogen followed by rubbing the leaf surface with a 9mm tapered arrow-end metal spatula. The collected trichomes were suspended in sterile water containing RNAprotect and vortexed for 30 seconds to dislodge the E. amylovora cells. RNA was isolated using the RNeasy Mini Kit (Qiagen, Germantown, MD, USA), and DNA was removed using the RNase-Free DNase Set (Qiagen, Germantown, MD, USA). The ‘inoculum’ sample underwent the same RNA isolation and purification procedure.

RNA processing and sequencing were performed at Yale Center for Genome Analysis (YCGA). RNA from the trichome samples underwent both bacterial and plant rRNA depletion procedures. Bacterial rRNA depletion was performed using the NEBNext rRNA Depletion Kit (Bacteria) (New England Biolabs, Ipswich, MA,USA). Plant rRNA depletion was performed using TruSeq Stranded Total RNA with Ribo-Zero Plant (Illumina, San Diego, CA, USA). RNA from the ‘Inoculum’ samples only underwent bacterial rRNA depletion. Sequencing was performed on an Illumina NovaSeq X Plus using a 25B flow cell, with paired-end 2×150bp reads at a depth of 5 million reads for the inoculum samples. Initially, the RNA from trichome samples was sequenced at 25 million reads, and was re-sequenced to generate 100-200 million more reads (for glandular trichome samples), and 600 million reads (for non-glandular trichome samples). Sequences from the two batches were combined for subsequent analysis and are available at the NCBI Sequence Read Archive PRJNA1166635.

Post-sequencing, reads were trimmed using Trimmomatic v0.39 (Bolger *et al*., 2014), and quality was assessed with FastQC v0.12.1(Andrews, 2010). The trimmed reads were aligned to the *Erwinia amylovora* ATCC 49946 reference genome using STAR v2.7.11b (Dobin *et al*., 2013). Alignment results indicated that there were 360-760 reads per gene, providing sufficient confidence for downstream analysis. Sequencing and alignment metrics can be found in Supplementary Table 1. Differential gene expression was analyzed using DESeq2 v3.19 (Love *et al*., 2014), and KEGG pathway analysis was conducted with BlastKOALA v3.0 (Kanehisa *et al*., 2016). Data visualization was performed using ggplot2, along with dplyr v1.1.2, readr v2.1.4, and cowplot v1.1.1 in RStudio. The codes for the bioinformatic analysis and data illustration can be accessed at https://github.com/quanzeng23/GlandularTrichomeLeafHair/tree/main.

## RESULTS

### *Erwinia amylovora* is able to infect apple shoots without artificial injuries

To determine whether *E. amylovora* can infect non-injured apple leaves, potted apple trees grown in a plant growth chamber free of any known mechanical (wind, rain or hailstorm) or insect injuries were dip-inoculated with a suspension of *E. amylovora* strain *Ea*110. Inoculated trees were maintained at 25 °C with 100% relative humidity and were monitored daily for disease symptoms daily. Initial symptoms appeared 2-3 days post inoculation (dpi), as necrosis on serrations on leaf margins and on veins (Supplementary Fig. 1A, B). Symptom progression was observed 4-5 dpi, as necrosis expanded from the tips of leaf serrations into internal leaf tissue, and along the midrib and veins. While infections of many veins seem to originate from the initial infection of nearby leaf margins, infection of some veins did not seem to have a clear linkage to a leaf margin, suggesting that leaf margins and veins could be independent entry points of the pathogen (Supplementary Fig. 1A). On day 5, 95% of the inoculated leaves showed shoot blight symptoms. These observations confirm that *E. amylovora* is able to infect apple leaves without artificial injuries.

### GUS staining identified GT and NT as potential epiphytic colonization sites and entry points

To identify epiphytic colonization sites and potential infection paths of *E. amylovora* on apple leaves, injury-free apple leaves were dip-inoculated in a suspension of *E. amylovora* strain *Ea*1189 with a *gus* (β-glucuronidase) gene integrated into its chromosome, and incubated under the same injury-free conditions described above. The presence of *E. amylovora* on apple leaves, indicated by the blue color after GUS staining, appeared as early as 17 hpi, on non-glandular trichomes (NTs) (Fig. 1A, red arrow) and glandular trichomes (GTs) (Fig. 1A, black arrow). For NTs, the presence of *E. amylovora* was more often detected on NTs near veins and midribs (Fig. 1B, left), and on leaf margins (Fig. 1B, right), than NTs between veins. For GTs, presence of *E. amylovora* was found on GTs located at the serrations of leaf margins (Fig. 1C, black arrow) and along the midribs and veins (Fig. 1C, red arrow). Thirty-six hours after inoculation, 0.8 ± 1.2% of the total NTs (± 1 mm adjacent to vein) and 4.3 ± 3.2% of the total GTs were colonized by *E. amylovora* as determined by GUS staining.

**Figure 1.**
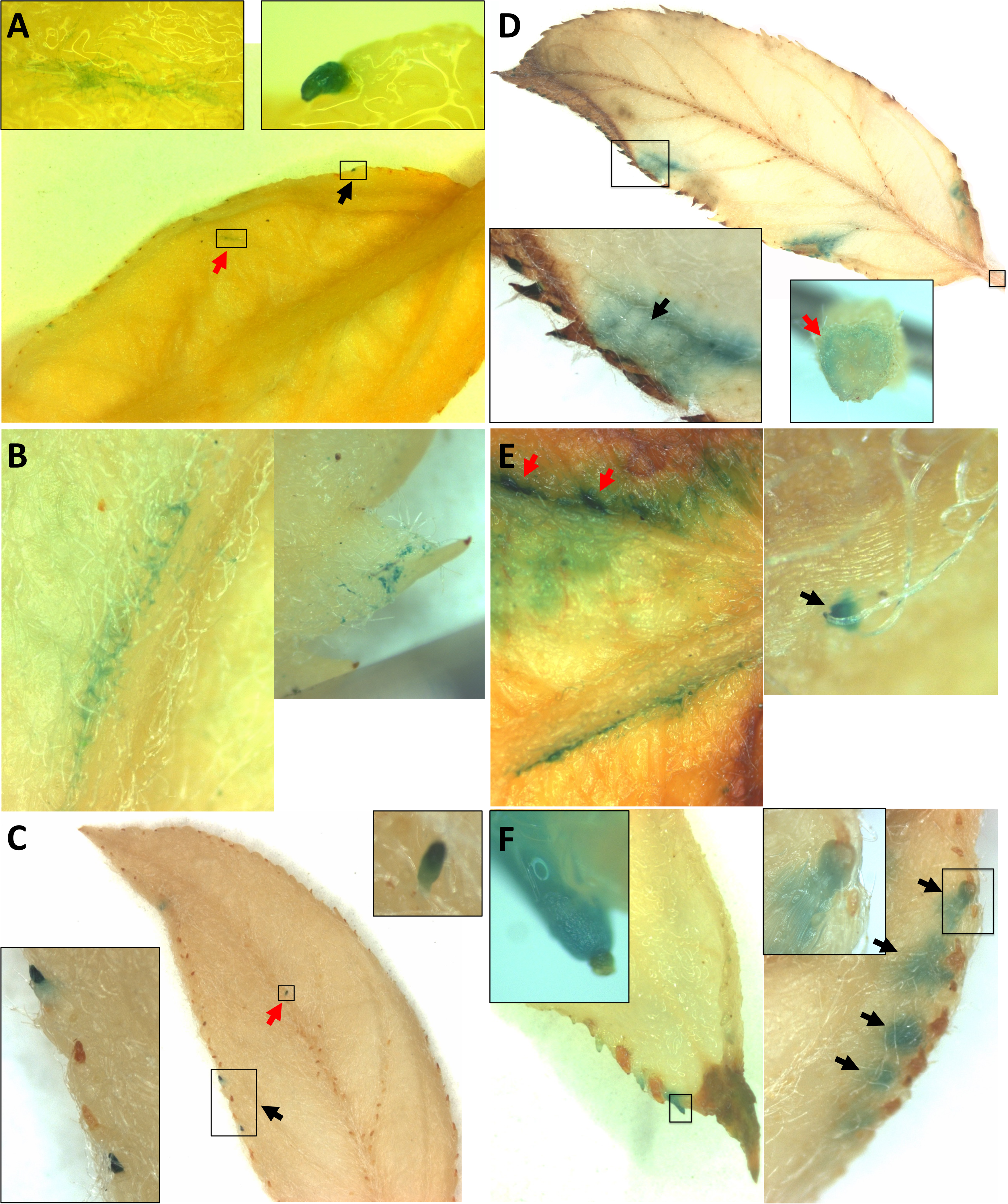
Detection of *E. amylovora* on apple leaves by GUS staining. Newly expanded apple leaves were inoculated with *E. amylovora*::*gus.* Samples were collected at different time points and processed for GUS staining. **A-C**, Detection of blue stain within the first 17 hr post inoculation (hpi) on NT (**A**, red arrow) and GT (**A**, black arrow). Blue stain on NTs was more frequently observed near the midvein (**B**, left) or near the margin (**B**, right), with little to no stain detected on NT in interveinal regions. Presence of blue stain on GTs located on both the margin (**C**, black arrow) and atop the midvein (**C**, red arrow), with blue stain on adjacent GTs seen to be random (**C** insert). **D-F**, Progressed infection of *E. amylovora* on dip-inoculated leaves (3-5 dpi). **D**, Image showing blue stain expanding from the initially infected GTs into the lateral vein (black arrow) and ultimately into the petiole (red arrow). **E**, Evidence showing colonization of *E. amylovora* on NTs can provide entry into leaf tissue (right) and can lead to vein infection (left). **F**, Evidence showing blue stain expanding from the infected GTs to the adjacent leaf tissue (black arrow).

Progressed colonization of *E. amylovora* was observed at 3-5 dpi, represented by both increased percentages of infected GTs and NTs, and expansion of *E. amylovora* presence into leaf epithem and veins (Fig. 1D). For NTs, the percentage of structures colonized by *E. amylovora* increased to 23.2 ± 13.5% at 3 dpi as indicated by GUS staining. Although the presence of *E. amylovora* could be found on NTs between veins, NTs near veins harbored a higher abundance of *E. amylovora* (Fig. 1E, red arrows). The presence of *E. amylovora* was often observed at the base of the NTs where they connect with the epidermal layer (Fig. 1E, black arrow).

For GTs, percentage of the GTs colonized by *E. amylovora* increased to 26.4 ± 9.8% at 3 dpi. Additionally, blue GUS stain expanded from the GTs to the apoplast in leaf serrations immediately connected to GTs (Fig. 1F). The location of blue stain in connected leaf tissue shows a matching pattern to the locations of the necrotic GTs (Fig. 1F, black arrows), which suggests the spread of *E. amylovora* from infected GT towards the internal leaf tissue. Eventually, *E. amylovora* infection reached minor veins (Fig. 1D, black arrow) and, subsequently, the leaf petiole (Fig. 1D, red arrow). Taken together, the GUS staining identified GTs and NTs as the epiphytic colonization sites of *E. amylovora* on apple leaves. It also depicts a potential entry and invasion from GTs and NTs to the connected leaf apoplast and ultimately xylem.

### Morphology of GTs and NTs on apple leaves

As the GTs and NTs are potential host entry points on apple leaves, their morphology and distribution on apple leaves were further examined. GTs are orange to brown colored, multicellular cortical structures that range from 0.09 to 0.20 mm in length and 0.04 to 0.11 mm in width (Supplementary Fig. 2A). GTs are located at the tips of serrations on leaf margins (Supplementary Fig. 2A), and on the adaxial leaf surface along the midvein and occasionally along the lateral veins (Fig. 1C). GTs on leaf serrations are located just beyond the termination of the vasculature at the edge of the leaf tissue (Supplementary Fig. 2A bottom left panel). Known as the epithem, this region is marked by loosely arranged parenchyma cells and large intercellular spaces (Supplementary Fig. 2A, right panel). No secretion outlets or any other natural openings were identified on the GT surface. Hydathodes can be found near the tip of the leaf serration adjacent to the GT within 75 to 110 μm (Supplementary Fig. 2A, right panel).

The NTs are hair-like structures attached to the epidermal layer on both adaxial and abaxial surfaces of the leaves, approximately 0.2 to 1.0 mm in length and 0.01 to 0.02 mm in width. NTs are more abundant on the abaxial surface than the adaxial surface (Supplementary Fig. 2B). On abaxial surface of younger leaves, NTs are highly abundant, forming a hairy layer covering the entire leaf surface (Supplementary Fig. 2B). No natural openings were found on NTs.

### Observation of the epiphytic colonization of *E. amylovora* on GTs and NTs

The macroscopic observation of *E. amylovora* presence on GTs and NTs through GUS staining prompted us to further visualize *E. amylovora* colonization on these structures using microscopy. Using confocal microscopy, we detected epiphytic colonization of a *gfp-*labeled *E. amylovora* on GTs and NTs. On GTs, *E. amylovora* cells were localized as aggregates on the surface of the GT and particularly near the junction of GT and leaf serrations (Fig. 2 A, B, and C). No aggregation or obvious evidence of entry was observed on the hydathodes near the tips of the leaf serration (Fig. 2D). In more progressed infection, *E. amylovora* cells were localized in the parenchyma tissue in the leaf serration near GTs (Fig. 2D). On NTs, epiphytic colonization of *E. amylovora* cells was also found on both stalks (Fig. 2E) and their base (Fig. 2F).

**Figure 2.**
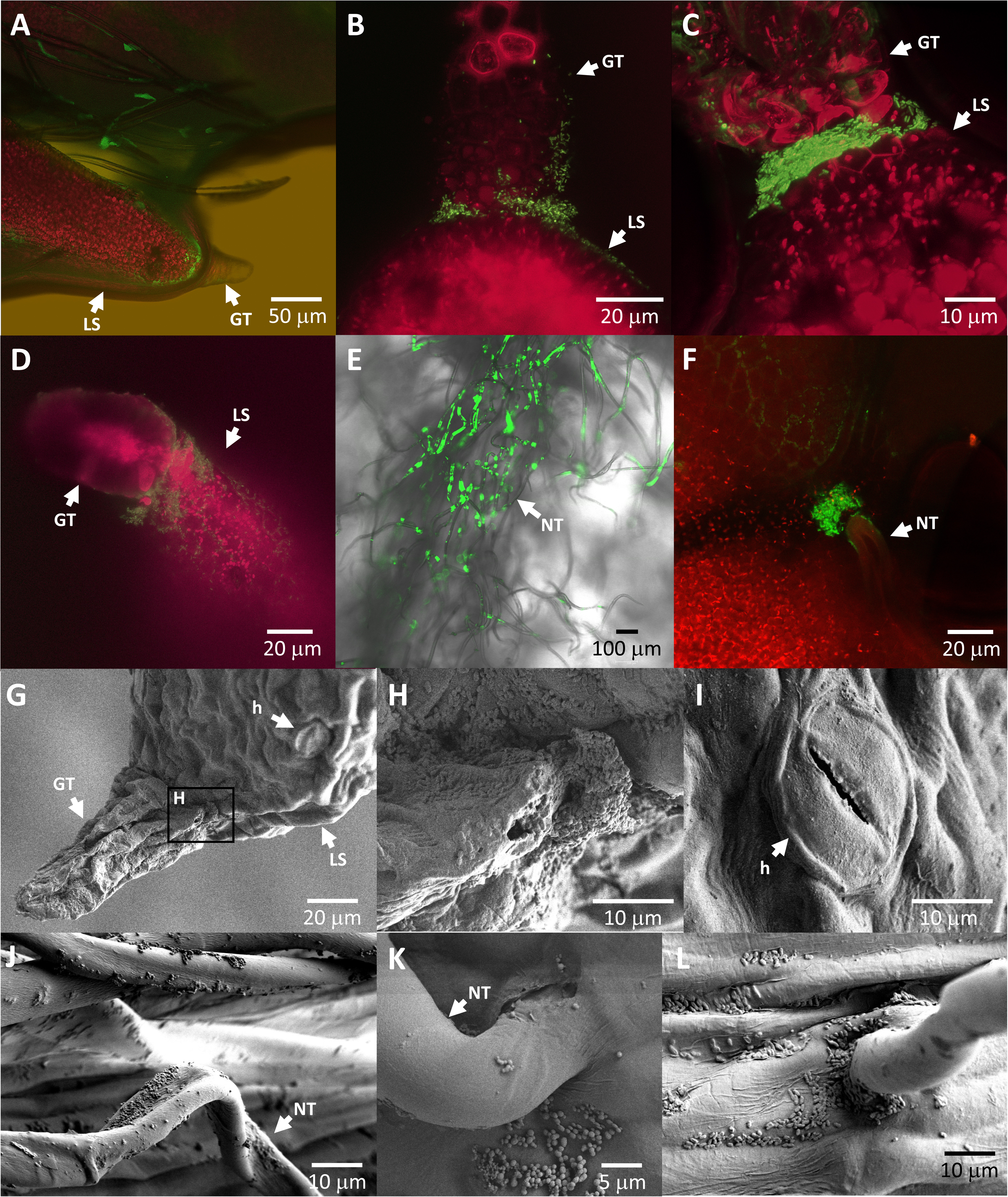
Detection of *E. amylovora* on apple leaves using confocal microscopy and scanning electron microscopy. **A-F**, Apple leaves dip inoculated in *E. amylovora*::*GFP* observed with a confocal microscope 3 dpi. *E. amylovora* is visualized as green fluorescent cells, autofluorescence of chloroplasts appears red. **A-D**, Large bacterial aggregates of *E. amylovora* on surface of GT and at the junction between GT and the leaf serration (LS), with few to no bacterial cells in LS further away from this junction. Some *E. amylovora* cells are visualized within the mesophyll tissue (**D**). Absence of *E. amylovora* cells on hydathode pores (h). **E-F**, *E. amylovora* on the surfaces of NT and the attachment point where a NT connects to the leaf epidermal layer. **G-L**, scanning electron micrographs of GT and NT at 3 dpi after dip inoculation with *E. amylovora*. **G-H**, *E. amylovora* cells are abundant at the junction between GT and a leaf serration (LS). **L**, no *E. amylovora* cells are seen at the hydathode pores (h) at the tip of the leaf serration. **J-L**, colonization of *E. amylovora* is observed on the surfaces of NT and at the base where they connect to the epidermal layer.

Epiphytic colonization of *E. amylovora* cells was also observed using scanning electron microscopy (SEM). Similar to the confocal microscopy observations, heavy *E. amylovora* colonization was observed within grooves and folds on the GT surface, and near the GT-leaf serration junction (Fig. 2, G and H), but not near hydathodes (Fig. 2I). *E. amylovora* cells were also found on NT stalk and base (Fig. 2 J, K, and L), and in grooves of leaf veins (Fig. 2L).

Similar to the GUS staining observation, no-significant colonization was observed on leaf surfaces lacking glandular and non-glandular trichomes. We also did not observe any evidence of bacterial aggregation near or entry through stomata.

### Quantification of epiphytic *E. amylovora* cells on GT and NT

To understand the epiphytic growth of *E. amylovora* on GT, we performed a temporal population analysis (10, 48, 72 hpi) on the GT of leaves dip-inoculated with *E. amylovora*. On average, 193 CFU of *E. amylovora* were present on each GT 10-hour post inoculation (hpi), which increased to 680 CFU / GT at 48 hpi and 1,724 at CFU / GT at 72 hpi (Fig. 3A). Only a small portion of those cells persisted and initiated epiphytic colonization on GT. Interestingly, the colonization of GT is highly uneven in distribution, even at 72 hpi, 33% of GT harbored 100 CFU or less per GT, while some GTs harbored as many as 9,250 CFU / GT.

**Figure 3.**
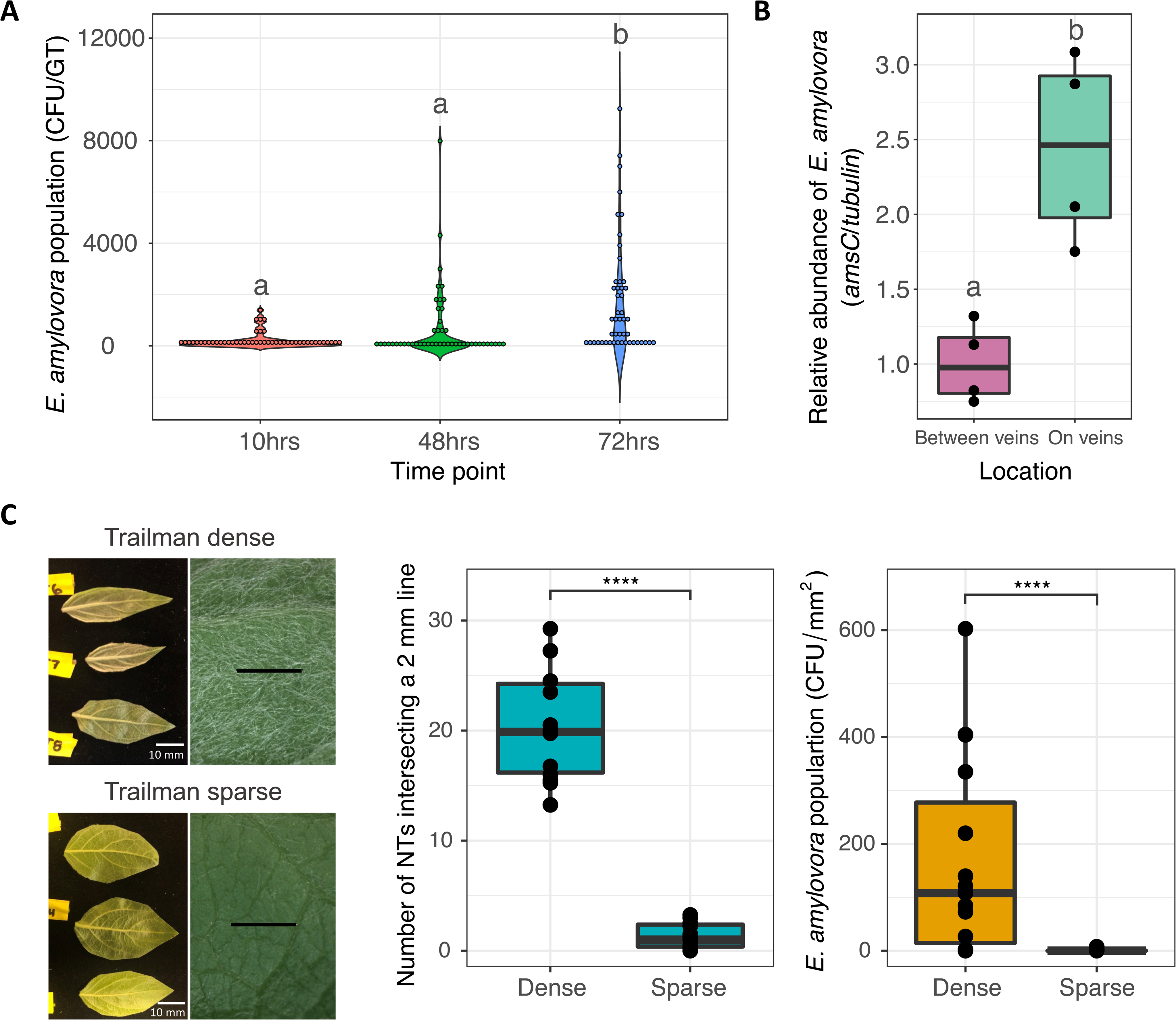
Colonization of *E. amylovora* on GT and NT. **A**, Population sizes of *E. amylovora* on GT at 10, 48, and 72 hpi, measured as colony forming units (CFU) per GT. Means separation lettering was determined using a nonparametric Kruskal-Wallis test (*P* =0.0041). **B**, Comparison of *E. amylovora* presence on NT collected from areas between the veins on the leaf and areas directly on the veins. *E. amylovora* presence, represented by the *amsC* copy number, was quantified by qPCR and was normalized by the copy number of the apple tubulin gene. **C**, Correlation of NT density and population size of *E. amylovora* on ‘Trailman’ leaves. Left: Morphology of newly developed leaves from apple cultivar ‘Trailman’ categorized as either “sparse” for low numbers of NT, or “dense” for high numbers of NT. Center: NT density was quantified by averaging four readings per leaf of the number of NT which intersected a 2 mm line. Statistical analysis was performed using a two-sample *t*-test (*P* < 0.0001). Right, Leaf wash was collected at 36 hpi with *E. amylovora* and was normalized as CFU/mm^2^ of leaf surface. The epiphytic population of *E. amylovora* was significantly higher at 168.27 CFU/mm^2^ on Trailman leaves in the dense category when compared with 0.6 CFU/mm^2^ from ‘Trailman’ leaves in the sparse category. Experiment was conducted three individual times, 5 leaf samples were collected from each cultivar per experiment. Statistical analysis was performed using a nonparametric Kruskal-Wallis test (*P* < 0.0001).

Due to the difficulty of manipulating the fine NTs, we used SEM images and estimated the epiphytic population of *E. amylovora* per NT. At 56 hpi, each NT harbored an average of 3.9 *E. amylovora* cells per NT, with some NT harboring as many as 782 CFU / NT. Using qPCR, we determined that NTs along veins harbored 2.5 times more *E. amylovora* cells than NTs between veins (Fig. 3B).

### Correlation of NT density with epiphytic population of *E. amylovora* on apple leaves

Under growth chamber conditions, we observed that the NT density of ‘Trailman’ apple leaves of the same age varied within a range (Fig. 3C), that provided an opportunity to determine a possible correlation between NT density with *E. amylovora* population on leaves. ‘Trailman’ leaves with high NT density (20.2 ± 4.9 NT per 2 mm) and with low NT density (1.3 ± 1.1 NT per 2 mm) were dip-inoculated with *E. amylovora*, and the epiphytic population of *E. amylovora* was determined at 36 hpi. As expected, leaves with higher NT density also displayed a significantly higher population of *E. amylovora* (168.3 cfu/mm^2^ leaf surface) as compared to leaves with lower NT density (0.6 cfu/mm^2^ leaf surface) (Fig. 3C). This observation further suggests that density of NTs significantly impact the epiphytic colonization of *E. amylovora*.

### Variation of NT density among different apple species and cultivars

Leaves from 34 different apple taxa, consisting of four *Malus* species (*M. domestica*, *M. bacatta*, *M. fusca*, and *M. sieversii*) and five *Malus* hybrids were analyzed for their NT density using a grayscale quantification method (Supplementary Fig. 3). The NT density of apple species and cultivars sampled ranged from 39.54 to 141.94 in their grayscale value (Supplementary Fig. 4). High density NT coverage seemed to be the normal condition among the varieties sampled, with the majority having a grayscale value higher than 75.0. Crabapples and *M. hybrid* varieties exhibited the lowest NT density.

### Evidence of *E. amylovora* entry into apple leaf through GTs

Using transmission electron microscopy, we examined the presence of *E. amylovora* in GTs and the connected epithem, during a course of infection. On leaves with no visible symptoms, epiphytic *E. amylovora* cells were found on grooves and folds on the GT, as well as at the junction of GT to the leaf serrations (Supplementary Fig. 5). Surface of leaf serrations farther away from GTs did not harbor any bacteria (Supplementary Fig. 5).

On leaves displaying necrosis at the tip of leaf serrations measuring < 1.0 mm^2^ revealed *E. amylovora* cells on the surface (Fig. 4B) as well as internally in plant tissue. *E. amylovora* cells were detected internally in GTs (Fig. 4C), at the junction between GTs and leaf serration (Fig. 4D), and inside the epithem (Fig. 4E). Within these locations, *E. amylovora* cells were found within the intercellular space between parenchyma cells, with large aggregation to spaces alongside the vasculature (Fig. 4E). Host cells in contact with *E. amylovora* appear highly degraded and shrunken (Fig. 4A, white arrows). These observations provide direct evidence that GTs are not only the epiphytic colonization sites but also entry points of *E. amylovora* during the infection of apple leaves.

**Figure 4.**
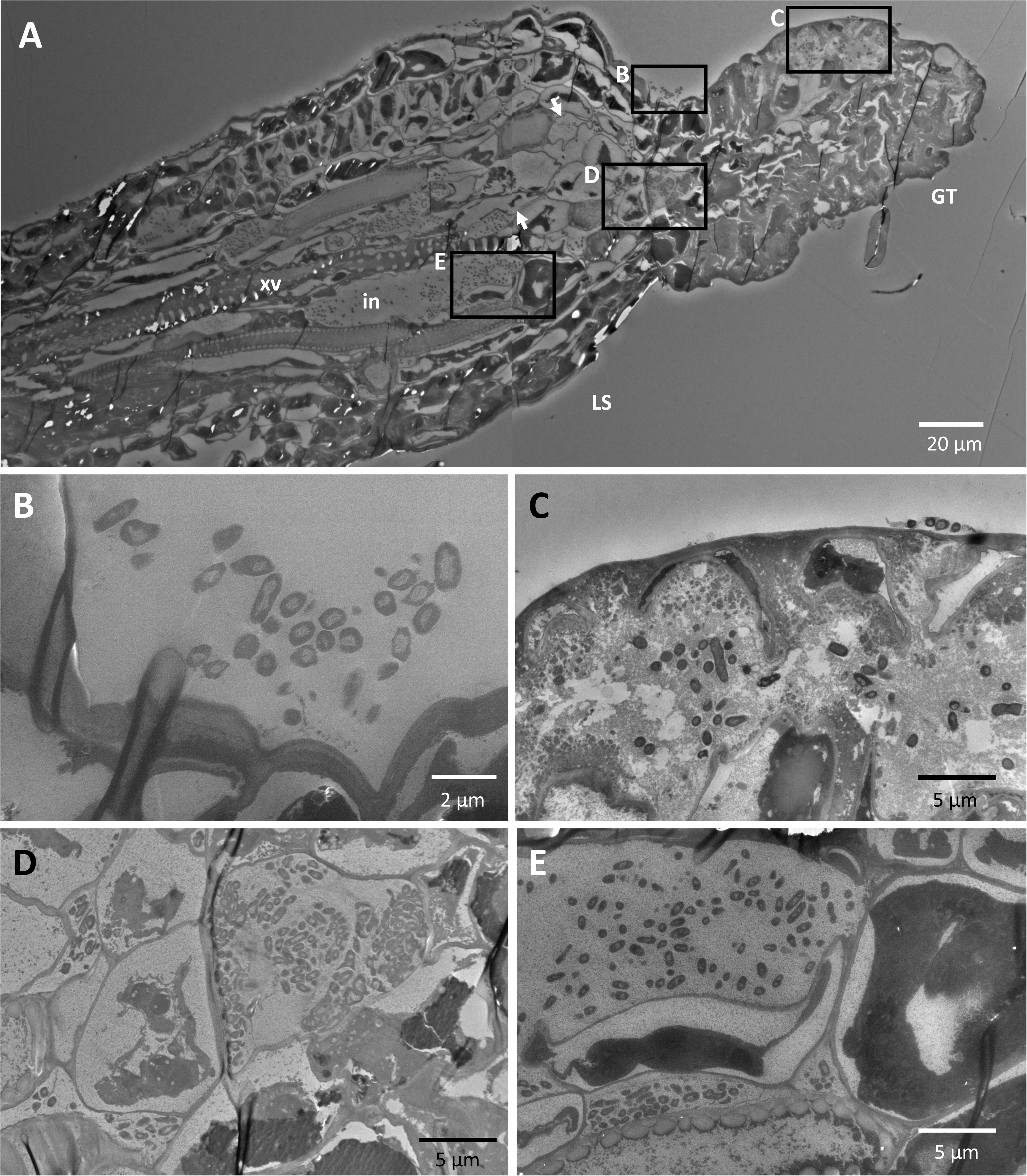
Observation of epiphytic and endophytic colonization of *E. amylovora* in GT by transmission electron microscopy. **A**, Single GT and tip of a leaf serration (LS) at 3 dpi showing progression of the bacteria into interior of the GT and the epithem region at the termination of the xylem vessels (xv). White arrows show shrunken host cells, pushed by intercellular pockets (in) of *E. amylovora.* **B-E**, Details of boxed areas in A. **B**, *E. amylovora* epiphytic colonization at the attachment point of the GT with the tip of the LS. **C**, Detail of *E. amylovora* cells within the GT. **D**, Detail of *E. amylovora* cells within the junction of the GT with the tip of the LS and leaf parenchyma cells exhibiting a shrinking of cell contents to the center of the cells. **E**, Detail of accumulation of *E. amylovora* cells within a pocket in one of several large intercellular spaces alongside xylem vessels (xv).

### Abscission of NTs and GTs during leaf ontogenesis

During leaf ontogenesis, it appeared that the abundance of NTs on the same leaf decreased over time. Mature apple leaves show far fewer NTs than the newly opened leaves (Fig. 5A). SEM images have shown ruptures at the base of NTs where they connect to leaf epidermis, and eventual abscission of the NTs (Fig. 5B, bottom panels).

**Figure 5.**
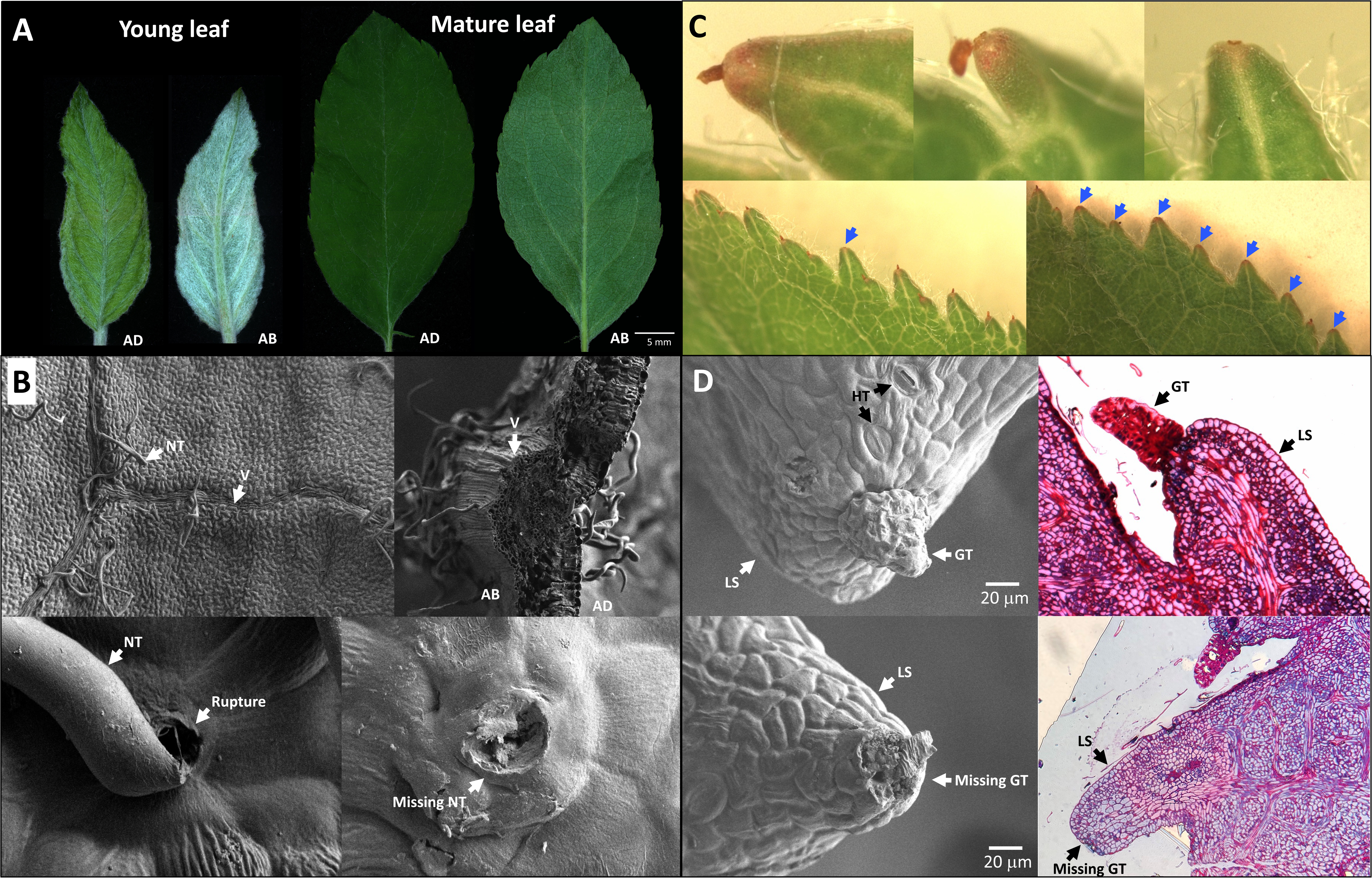
Abscission of GT and NT during the course of leaf development. **A**, Young apple leaves (left) have more NTs when they first unfold compared with mature apple leaves (right) from the same tree several weeks after unfolding. Grayscale values representing the NT density for the adaxial and abaxial surfaces were 80.76 and 145.08 respectively for the young leaf, and 32.09 and 64.40 respectively for the mature leaf. **B**, Scanning electron micrographs of the connection of NT to the abaxial leaf epidermal layer (top left) and on both surfaces in cross section (top right), as well as ruptures which can occur at their base (bottom left) before the eventual loss of a NT (bottom right). **C**, Young leaves (bottom left) have GTs attached to the majority of their LS during the first two three weeks after unfolding. Nearly all GT on LS are missing in mature leaves (blue arrows, bottom right). **D**, Scanning electron micrograph and semi-thin section of leaf serrations from young leaves (top row) showing intact GT. Scanning electron micrograph and semi-thin section of leaf serrations from mature leaves (bottom row) showing missing GT and abscission point.

Similar to NTs, abscission of GTs related to leaf ontogeny was also observed (Fig. 5C, D and Fig. 6B beige bar). Partly unfolded young apple leaves had GT attached to the tips of all of their leaf serrations (stage 1 in Fig. 6B). The breakoff of GTs was initiated approximately 4 to 5 days after leaf unfolding (stage 2 in Fig. 6B). By 10 to 14 days after unfolding, 46% of leaf serrations lost the GT (stage 4 in Fig. 6B). By 28 days, only 22% of leaves had GTs still attached at the tips of leaf serrations (stage 6 in Fig. 6B). This suggests that GTs naturally abscised over the course of leaf expansion and maturation, independent of any external disturbance in the growth chamber condition.

**Figure 6.**
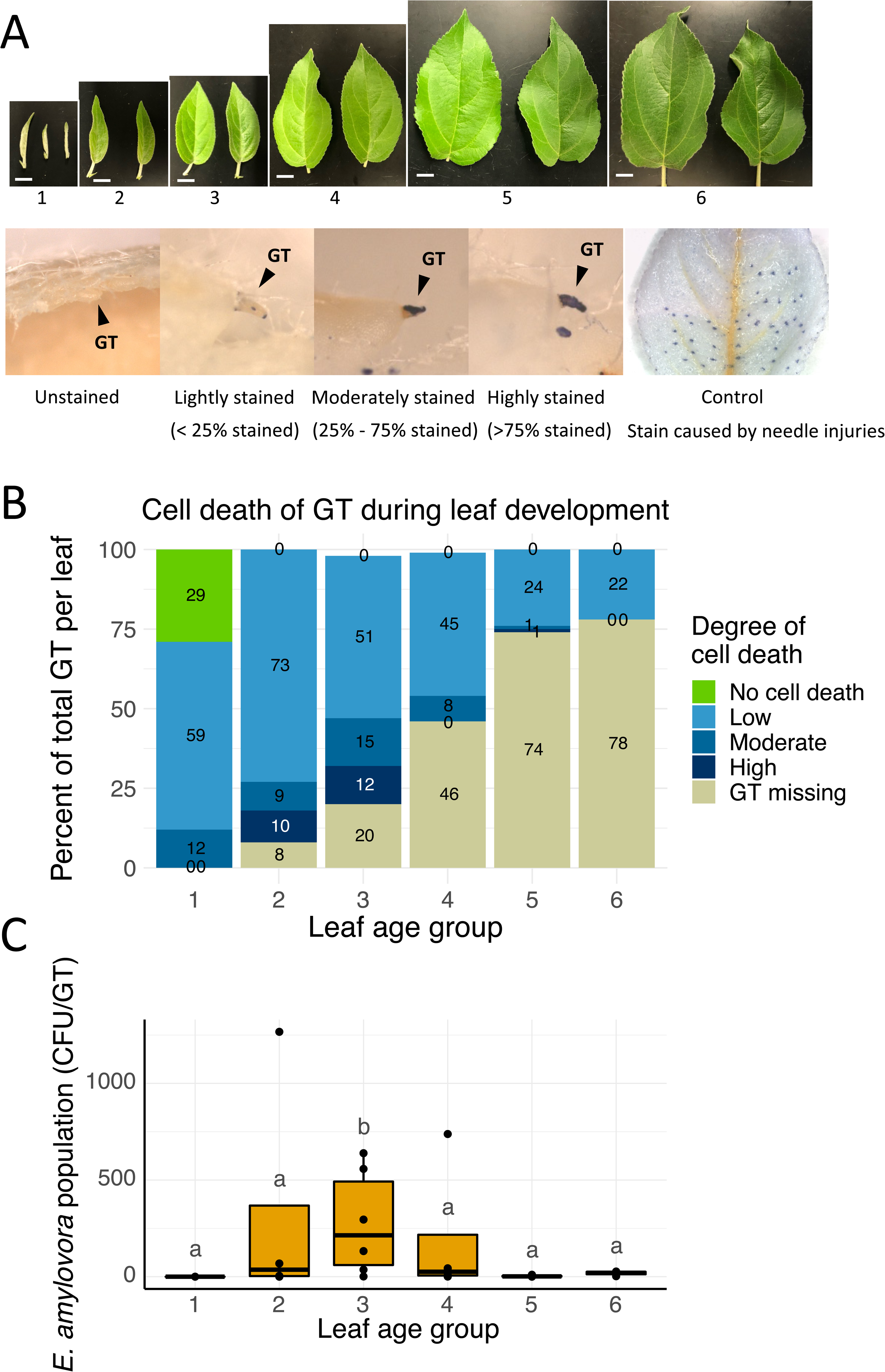
Observations of wounding and loss of GT during leaf development and *E. amylovora* populations on leaves of different ages. **A**, Representative images of leaves from the ‘Frostbite’ cultivar showing 6 age groups and different levels of GT injuries. Top panel: leaf development stages defined by the number of days past leaf unfolding (dpu): stage 1: unopened leaf; stage 2: 4 or 5 dpu; stage 3: 5-10 dpu; stage 4: 10 - 14 dpu, stage 5: 14 - 28 dpu, stage 6: greater than 28 dpu. Bottom panel: wound severity divided into 4 categories based on percent of GT surface area that are blue in trypan blue staining: unstained GT (indicating no cell death), Low (< 25% GT surface stained), Moderate (between 25% and 75% GT surface stained), and High (> 75% GT surface stained). Leaves artificially injured by needle poking showing blue stains at the injury points were used as controls. **B**, Analysis of the severity of wounding on GT from each of 6 age groups. **C**, Population of *E. amylovora* on dip inoculated leaves from each of 6 age groups at 3.5 dpi. Population size was reported as CFU per GT. Experiment was performed twice, with between 18 and 25 GT collected from each sample, between 2 and 5 leaves per leaf stage in each experiment and similar results observed.

### Naturally-occurring host cell death of GTs during leaf development is essential for *E. amylovora* colonization

As the microscopy examinations did not identify any secretion outlets or natural openings on GTs, we next examined whether any naturally-occurring wounds are present on GTs during the abscission process, and whether such wounds are beneficial to *E. amylovora* infection. Using trypan blue staining, wounded cells (cell death) could be differentiated from healthy cells by the blue color (Fig. 6A). Using this method, the presence of cell death on GTs was examined over a period of leaf development (from leaf unfolding to mature leaf, over 4 weeks; stages 1-6 in Fig. 6A). Level of cell death on an individual GT is categorized into low, moderate, or high based on the amount of staining observed (Fig. 6A). Within stage 1 leaves, 29% of GTs exhibited no cell death, while the remaining 71% showed low and moderate degree of cell death (Fig. 6B). No abscission of GTs was observed on stage 1 leaves. Abscission of GTs was first observed on stage 2 leaves, about 4 – 5 days after the leaf fully unfolded. Co-occurring with the abscission, increase of plant cell death was observed on GTs during the same period, with 19% and 27% of GTs displaying at least a moderate level of cell death on stage 2 and stage 3 leaves (Fig. 6B).

While the abscission of GTs continued into leaf stages 4-6, a decline in cell death on GTs was observed once leaves reached stage 4, with only 8% of GTs exhibiting moderate cell death, and none of them showing a high level of cell death. On mature leaves (stage 5 and 6), almost none of the GTs displayed any moderate or high level of cell death (Fig 6B).

To determine whether the naturally-occurring cell death is associated with *E. amylovora* colonization on GTs, pathogen load of *E. amylovora* on GTs was quantified in tandem with the level of cell death during leaf development (Fig. 6C). Importantly, the *E. amylovora* population followed the same trend as the presence of host cell death: higher populations of *E. amylovora* cells were recovered from stage 2 and stage 3 leaves. Partially unfolded leaves with GTs present but lack of cell death (stage 1), or mature leaves with GTs missing and with limited cell death (stages 5 and 6) did not support *E. amylovora* growth at those structures. This suggests that naturally occurring wounding on GTs during leaf development is essential for *E. amylovora* colonization.

### Identification of differentially expressed genes (DEGs) in *E. amylovora* during colonization of GT and NT

To identify genes required for GT and NT colonization and the subsequent leaf invasion, a transcriptomic analysis was performed to search for differentially expressed genes (DEGs) in *E. amylovora* three days after inoculation on GT and NTs as compared to the inoculum. On GTs, 1,082 DEGs were identified in *E. amylovora*, with 553 upregulated and 529 down regulated genes (Supplementary table 2, Fig. 7A). The DEGs were categorized by their KEGG pathway functions (Fig. 7B). KEGG pathways with more upregulated DEGs than downregulated DEGs include transporters (Up: 94/Down:46), ribosome (Up: 35/Down:0), bacterial motility (Up: 34/Down:6), secretion system (Up: 26/Down:8), DNA repair and recombination (Up: 19/Down:7), lipopolysaccharide biosynthesis (Up:11/Down:3), amylovoran biosynthesis (Up: 11/Down:0), and peptidoglycan biosynthesis and degradation (Up: 11/Down:3). In contrast, pathways dominated by downregulated DEGs include the prokaryotic defense system (Up: 2/Down: 10), unknown functions (Up: 7/Down: 17), and enzymes with EC numbers (Up: 14/Down: 21) (Supplementary Table 2, Fig. 7B).

**Figure 7.**
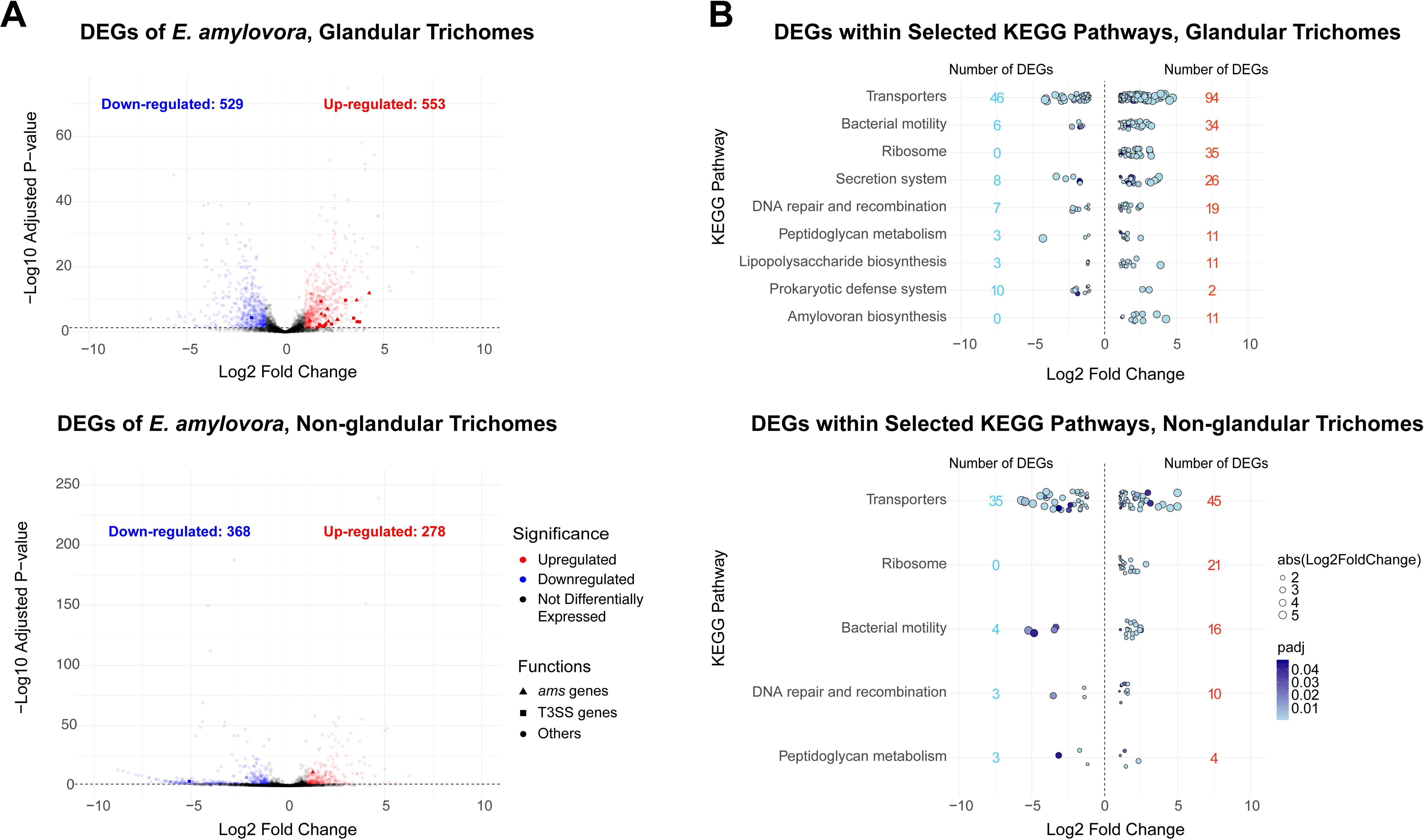
Differentially expressed genes (DEG) in *E. amylovora* during colonization of glandular and non-glandular trichomes. **A,** Volcano plot of DEGs identified in *E. amylovora.* **B,** Number of DEGs identified in each selected KEGG pathways.

Given that secretion systems are among the upregulated KEGG pathways in GTs, and that the type III secretion system (T3SS) is a critical virulence factor in *E. amylovora*, we examined the expression of T3SS genes more closely (Fig. 7A). Notably, 14 of the 40 known T3SS structural and effector genes are upregulated in GTs. Furthermore, genes involved in amylovoran biosynthesis, another key virulence factor, were also significantly upregulated during GT colonization, with 11 of the 12 known genes in this pathway showing upregulation (Fig. 7A).

On NTs, 646 DEGs were identified in *E. amylovora*, including 278 upregulated and 368 downregulated genes (Supplementary Table 2, Fig. 7A). Upregulated functions on NTs include bacterial motility (Up: 16/Down: 4), DNA repair and recombination (Up: 10/Down: 3), and ribosomal activity (Up: 21/Down: 0). Downregulated genes are associated with functions such as enzymes with EC numbers (Up: 9/Down: 11), peptidases and inhibitors (Up: 3/Down: 7), and transfer RNA biogenesis (Up: 3/Down: 9) (Supplementary Table 2, Fig. 7B).

### Contribution of the type III secretion system and amylovoran biosynthesis during colonization and entry of GT

To validate whether the T3SS and amylovoran biosynthesis genes were important for GT colonization and entry, deletion mutants of *hrpL*, the master regulator of the T3SS and the *ams* operon (Zhao *et al*., 2009) were inserted with the *gus* gene in their chromosome and were tested for their colonization on GTs during dip inoculation. Colonization of the GTs was not affected by the mutation of *ams* but was significantly reduced by mutation of *hrpL* (Fig. 8). While the *ams* mutant was still able to infect the epithem of some GTs, we did not observe any systemic spread from there to the nearby veins. The *hrpL* mutant was unable to cause any infection in the epithem (Fig. 8).

**Figure 8.**
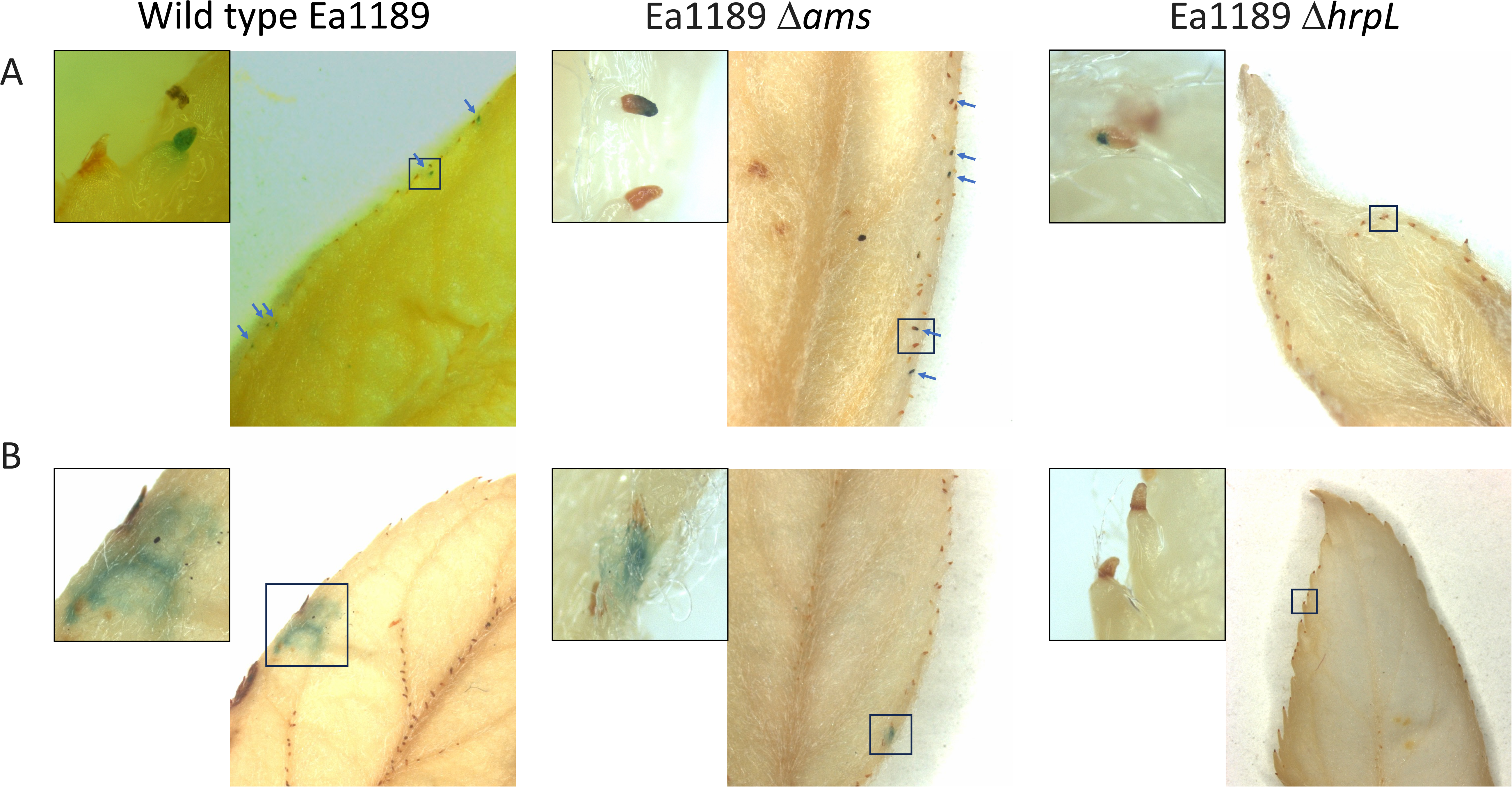
Colonization and infection of *E. amylovora* wild type, Δ*hrpL* and Δ*ams* on glandular trichomes of apple leaves. **A**, Initial infection stage (2 dpi); **B,** progressed infection stage (5 dpi).

## DISCUSSION

In this study, we report a novel mechanism utilized by bacterial plant pathogens to overcome the plant epidermis and enter their host. Added to the previously known entry mechanisms which include through natural openings (e.g. stomata and hydathodes) and mechanical injuries (e.g. those caused by feeding of piecing-sucking insects and by extreme weather events), we demonstrated that plant pathogenic bacteria may take advantage of injuries that occur during plant development such as abscission and enter hosts.

This discovery was made possible by the usage of a combined methods of macroscopic and microscopic observations. The macroscopic observation allows the identification of the epiphytic colonization sites as well as invasion paths at the whole leaf level through naked eyes. This not only identified GTs and NTs as the colonization sites of the pathogen, but also excluded other possible entries points such as stomata. NTs have been previously identified as a colonization sites for *E. amylovora* using fluorescent microscopy (Bogs et al., 1998). Through the macroscopic observation and later confirmed with qPCR, we observed that NTs located on veins harbor more *E. amylovora* cells than NTs located on between-vein leaf surface, which leads to infection of those veins. Importantly, the macroscopic observation of leaves at different infection stages provided evidence that the colonization of GTs and NTs by *E. amylovora* eventually led to infection in the nearby apoplast tissue and spread to the plant vasculature. As a complement to the macroscopic observations, our microscopic observations depict the *E. amylovora* cellcolonization pattern on GTs and NTs, and the evidence of an endophytic presence of *E. amylovora* at the junction of GTs to the leaf serration. Together, we depict a clear picture that *E. amylovora* can colonize GTs and NTs epiphytically, and enter hosts through GTs at leaf serrations and NTs on veins and cause shoot blight infection on apple leaves.

Trichomes are the most noticeable epidermal outgrowth on plant surfaces. These structures not only create complex three-dimensional structures, but also secrete secondary metabolites. Function-wise, trichomes contribute to the reduction in water loss, protection against herbivores, and regulation of temperature. However, their roles in plant-pathogen interactions and plant disease infection are poorly understood. Previous reports have shown that these structures are genotype-specific microbial hotspots in the tomato phyllosphere (Kusstatscher *et al*., 2020). Here, we showed that trichomes are also critical colonization sites and entry points of a plant pathogenic bacterium. We further showed that leaves with higher NT abundance harbor more *E. amylovora* cells than leaves with less NTs, and that different *Malus* species display a wide range of NT density. In this regard, GTs and NTs could be potential plant traits that need to be considered during disease resistance breeding.

We describe a reduction in trichome density and integrity on *Malus domestica* leaves during their development and maturation. As GTs and NTs are important colonization sites of *E. amylovora*, our findings provided partial explanation to a previous observation in the field that fire blight infection always initiates on young tender shoots rather than on mature, leathery leaves. More interestingly, using trypan blue staining, naturally occurred injuries were detected on GTs during this process, which is highly correlated with the *E. amylovora* quantification.

Changes in trichome abundance are probably related to the higher need for prevention of water loss, protection of UV and herbivore damage on young leaves than on mature leaves. However, a cost of this is the potential entry of pathogens through the wounds formed during this process. It is important to note that the naturally occurred injuries during this process, but not the abscission itself, are important for pathogen entry, as we showed that *E. amylovora* enters through GTs while GTs are still attached to the leaf serrations. We think that the abscission junctions callus during the abscission process and are no longer susceptible once GTs are lost. The entry through naturally occurred injuries is not restricted to leaf infection, as we also observed that *E. amylovora* can infect flowers through wounds created by petal abscission points during flower development (Data not shown).

Through transcriptomic analysis, our study identified key DEGs crucial for *E. amylovora* colonization of GTs and NTs. Notably, we demonstrated that two major virulence factors—the T3SS and amylovoran biosynthesis—are significantly upregulated in *E. amylovora* on GTs. This finding is consistent with previous studies, where T3SS and amylovoran-related genes were also shown to be induced during colonization of the stigma and fruitlet (Zhao et al., 2005; Schachterle et al., 2022). The importance of these genes in GT colonization on apple leaves was further validated through mutation, inoculation, and GUS staining experiments. These experiments revealed that mutations in these genes significantly impaired *E. amylovora*’s ability to colonize GTs.

In comparing the DEGs identified during GT and NT colonization, we highlight both shared and distinct functional expression patterns between the two sites. Ribosomal genes are upregulated in *E. amylovora* on both GTs and NTs, consistent with the known relationship between ribosome copy counts and cellular metabolic activity (Fu & Gong, 2017). This suggests that colonizing plant surface structures may elevate *E. amylovora*’s metabolic activity. Similarly, motility genes are upregulated at both sites, mirroring observations from flower infection (Schachterle *et al*., 2022). Given that *E. amylovora* enters hosts through naturally occurring wounds, increased motility likely facilitates more efficient movement towards these entry points. In contrast, transporter genes show higher upregulation on GTs (Up: 94/Down: 46) than on NTs (Up: 45/Down: 35). This may be due to the fact that GTs, but not NTs, actively secrete secondary metabolites. An enhanced transporter system could help *E. amylovora* more effectively utilize nutrients from GT secretions, supporting bacterial growth and establishing a larger population for infection.

Finally, findings from this study provide a much-needed answer as to how shoot blight is initiated in the orchard, and provided important implications to developing management strategies against fire blight. Most extension publications identified mechanical injuries, such as ones caused by wind gusts, blowing sand, hailstorms as well as piecing-sucking insects as the main driver for initiation of shoot blight infection (Johnson, 2015). Despite this, the presence of these micro-injuries have never been proven experimentally, and growers often observe shoot blight infection in their orchards regardless of any extreme weather events / significant insect activities. Some orchard owners planted wind blocking trees around orchard perimeters to reduce the impact of gusty winds and blowing sand. While these environmental factors and insect activities are indeed important in spreading the pathogen and may create injuries to shoots, our study showed that *E. amylovora* entry into shoots can happen naturally independent of those environmental stressors. Shoot blight often occurs in the field in a sporadic manner although there is a lack of understanding of the sporadic nature. Our study showed that even in full contact with a high population of *E. amylovora*, GTs on the same leaf harbor a wide range of *E. amylovora* cells, ranging from 10 to 10^4^ CFU per GT, which is about 10^2^-10^5^ times lower than the population found on stigmas of flowers (Malnoy *et al*., 2012; Slack *et al*., 2022). These findings suggests that the sporadic colonization and entry through GTs could be one explanation for the sporadic occurrence of the shoot blight infection.

## ACKNOWLEDGEMENTS

We would like to thank Jonathan Jacobs for his mentorship to Felicia Millett and suggestions during the research investigation and thesis drafting. We thank Maria Jose Estrada for her assistance in isolating RNAs from trichome samples; Antariksh Tyagi for assistance in assistance in RNAseq methods; and M. Amine Hassani for helping with figure illustrations. This research is supported by United State Department of Agriculture Specialty Crop Research Initiative (2023-51181-41319), Organic Agriculture Research and Extension Initiative (2023-51300-40727), Michigan State University AgBioResearch.

## AUTHOR CONTRIBUTIONS

QZ, FM, and James S designed the research. FM, James S, Jules S, KM, NZ, MA, performed the experiment. FM, James S, YL analyzed the data.JI, VR, GS, YL and QZ provided experimental resources, project management, and mentorship to the students. FM, James S, and QZ wrote the manuscript. All authors edited and approved the manuscripts.

## DATA AVAILABILITY

All sequencing reads used in this study are available at the NCBI Sequence Read Archive BioProject number PRJNA1166635. The codes for the bioinformatic analysis and data illustration can be accessed at https://github.com/quanzeng23/GlandularTrichomeLeafHair/tree/main.

## FIGURE LEGENDS

**Supplementary Figure 1.** Fire blight symptom on dip-inoculated apple leaves without artificial injury. **A**, Initial symptoms were most commonly observed on the leaf margins (left) and occasionally observed on the midvein or lateral veins (right). **B**, Symptom development over the course of 5 days post inoculation (dpi). Progression of symptoms typically begins on leaf margins and spreads to the midvein. Newly expanded apple leaves of two-year-old potted apple trees cultivar ‘Gala’ were dipped in a bacterial suspension containing 10^8^ CFU/ml of *E. amylovora*. Inoculated trees were maintained in a plant growth chamber that is free of insects and any other mechanical damage from the environment.

**Supplementary Figure 2.** Morphology of GT (**A**) and NT (**B**) on apple leaves. **A**, Morphology of GT on ‘Gala’ apple leaf margin examined by light microscopy (left panels) and by transmission electron microscopy (right panel). Within the epithem of the leaf, xylem vessels (xv) are identifiable by their secondary cell wall thickening. Loosely arranged parenchyma cells (p) accompany the vasculature and are separated by large intercellular spaces (in). The outer layer consists of epidermal cells (e), with an occasional hydathode pore (h). **B**, Presence of NT on abaxial and adaxial surfaces of ‘Pumpkin Russet’ apple leaves (top left and bottom). The abaxial surface (AB) has a higher density of NT than the adaxial surface (AD). Areas near the midvein and lateral veins (LV) have higher densities of NT than areas between vein surfaces. Transmission electron microscopy observation of a NT attached to the epidermal layer (top right). Long, ribbon shaped NT do not lay flat and often appear in cross section rather than longitudinally in TEM.

**Supplementary Figure 3.** Workflow for measuring pixel grayscale value from leaf images in ImageJ.

**Supplementary Figure 4.** Analysis of NT density across 34 apple taxa using pixel grayscale value as an indicator of trichome density. **A**, Representative images of leaves whose grayscale values placed the lowest, the highest, and one from middle of the range, with their grayscale values averaging 39.54, 96.8, and 141.94 respectively. **B**, Leaf samples from across 34 taxa including 4 different apple species, 5 hybrids, and 25 cultivars of *M. domestica* were evaluated for NT density on young leaves. Measurements are the average grayscale value of pixels from the abaxial and adaxial surfaces of each individual leaf. Four to six biological replicates were included in each taxa. Higher values are indicative of a greater abundance of NT covering the leaf surface as they reflect more light. The pixel grayscale value ranges from 0 (black) to 255 (white).

**Supplementary Figure 5.** Observation of *E. amylovora* epiphytic colonization of a GT by transmission electron microscopy. **A**, Epiphytic colonization of a GT at 3 dpi. The interior of the adjacent leaf serration (LS) does not contain bacterial cells. **B-D**, Details of *E. amylovora* cells along cavities on GT surface.

**Supplementary Table 1.** Metrics of RNAseq and alignment.

**Supplementary Table 2.** List of the differentially expressed genes (DEGs) identified in E. amylovora during colonization of glandular and non-glandular trichomes.

