## Supplementary figures and images for "The fire blight pathogen *Erwinia amylovora* enters apple leaves through naturally-occurring wounds from the abscission of trichomes"

### Supplementary Fig. 1

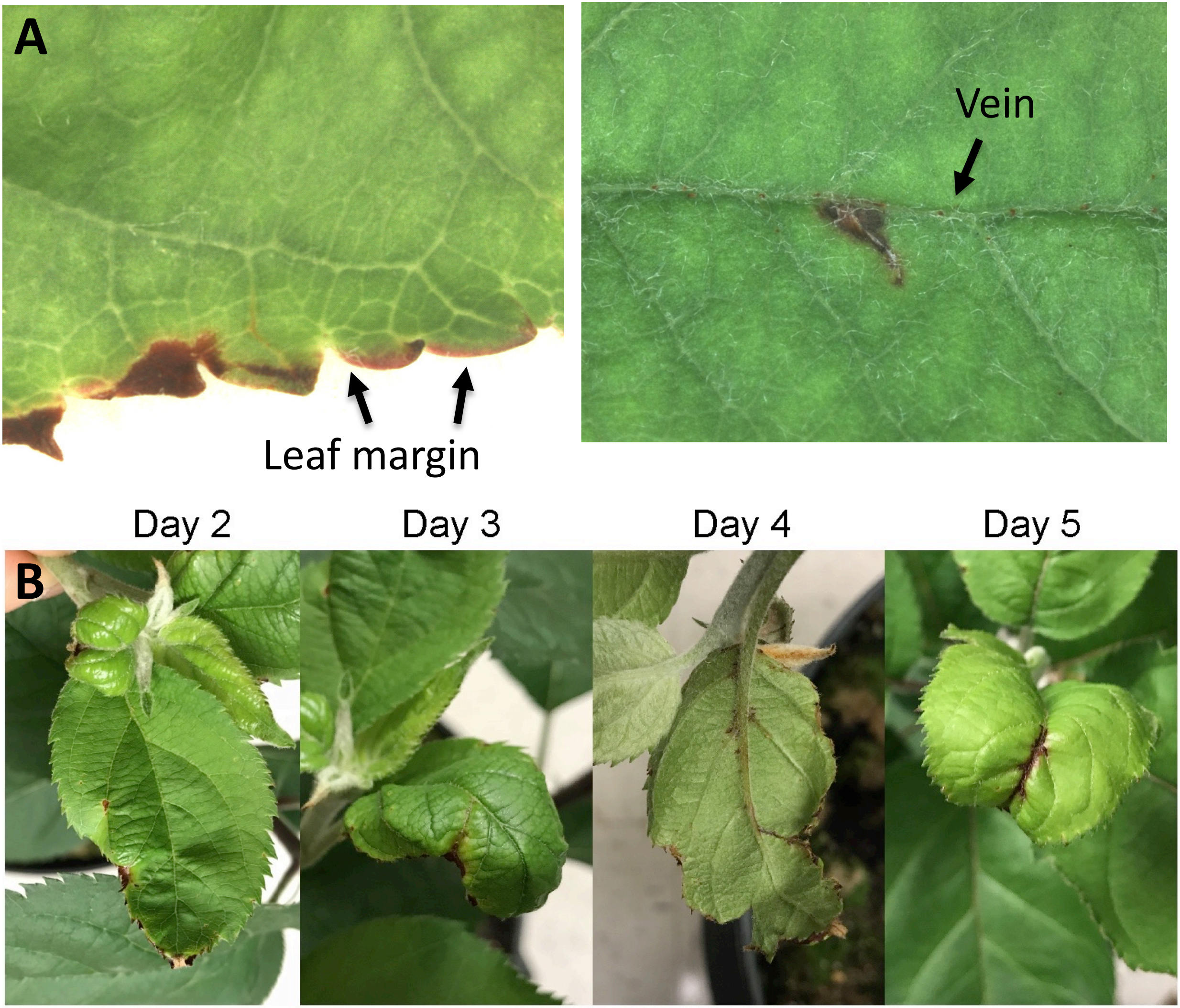

### Supplementary Fig. 2

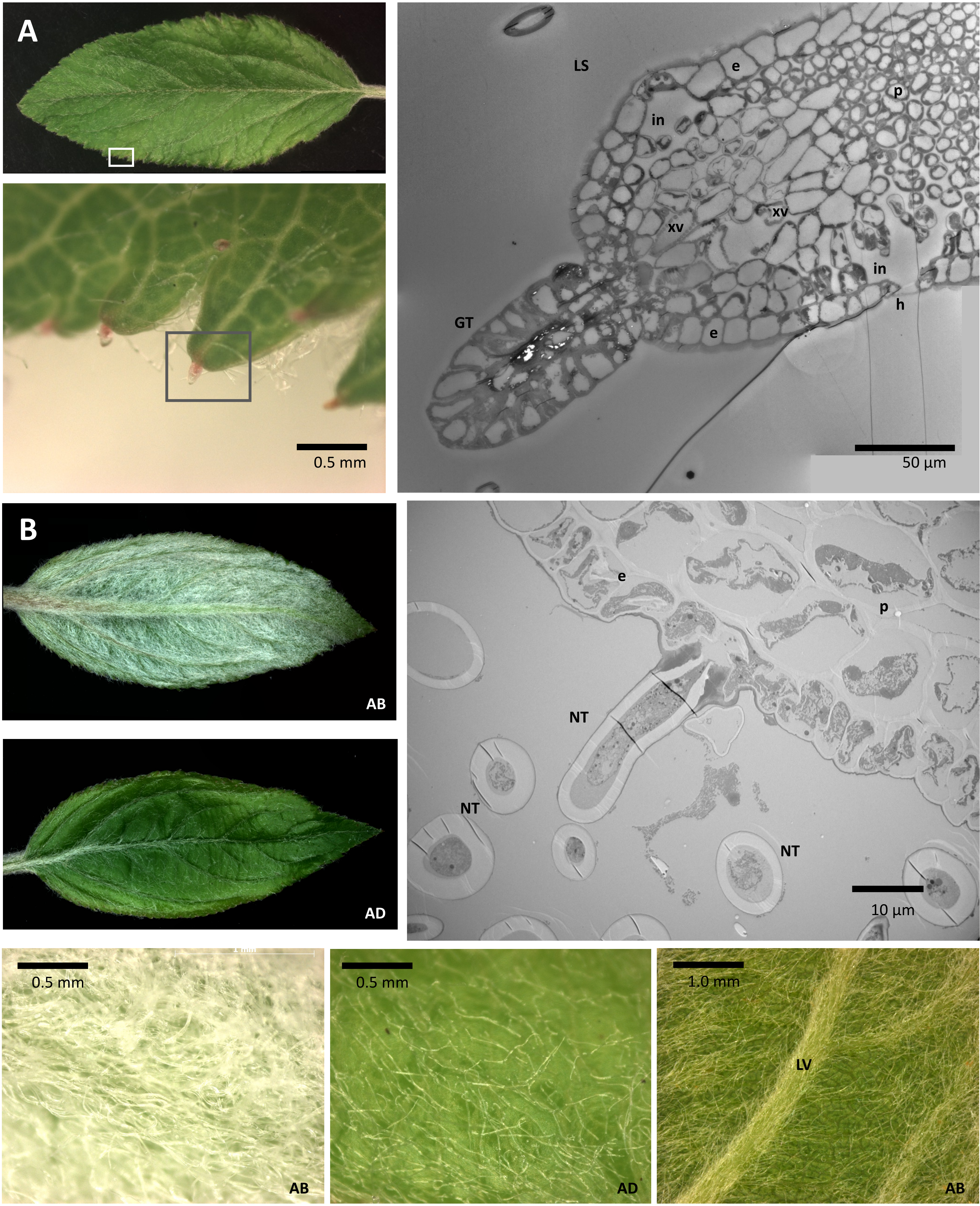

### Supplementary Fig. 3

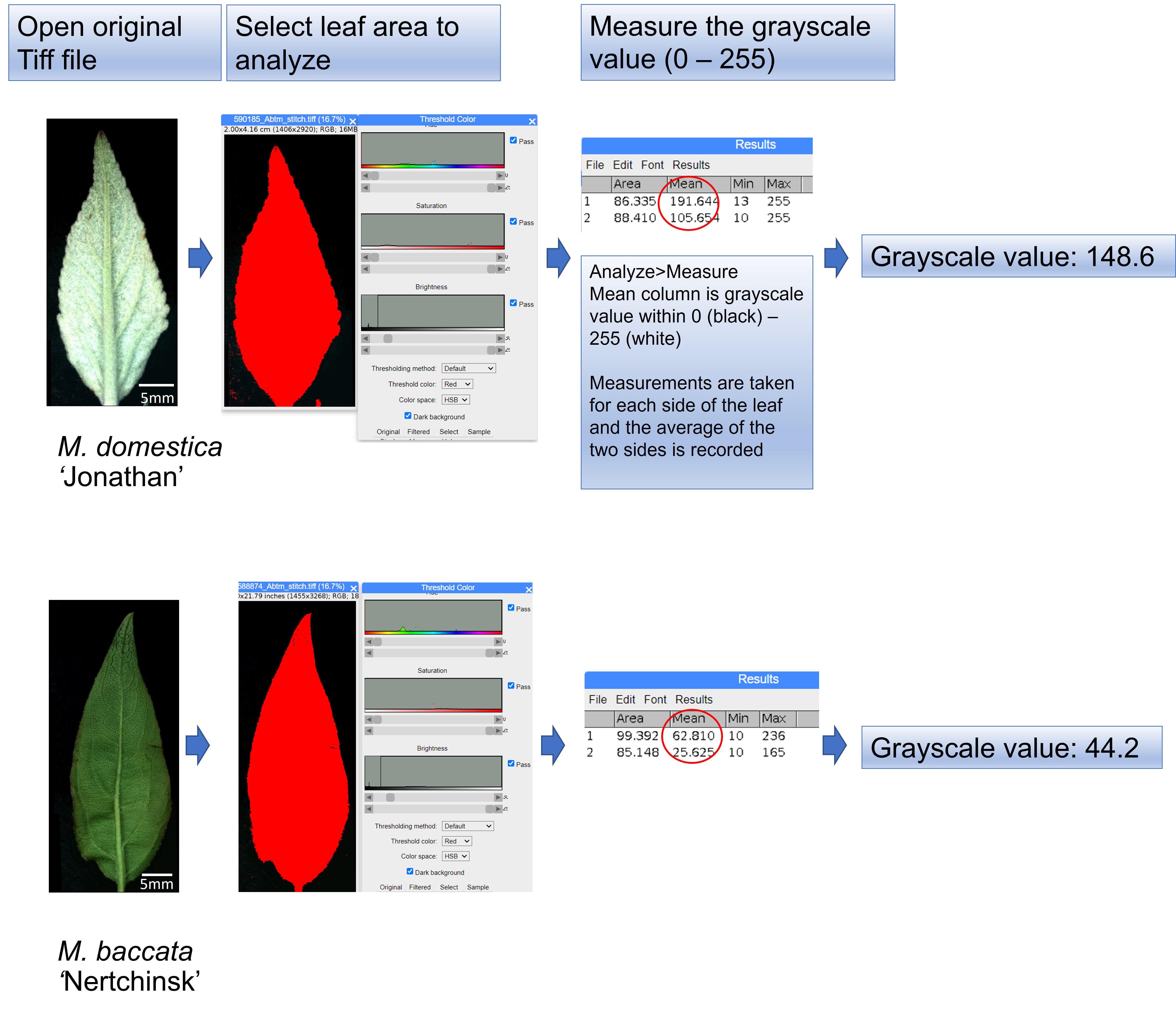

### Supplementary Fig. 4

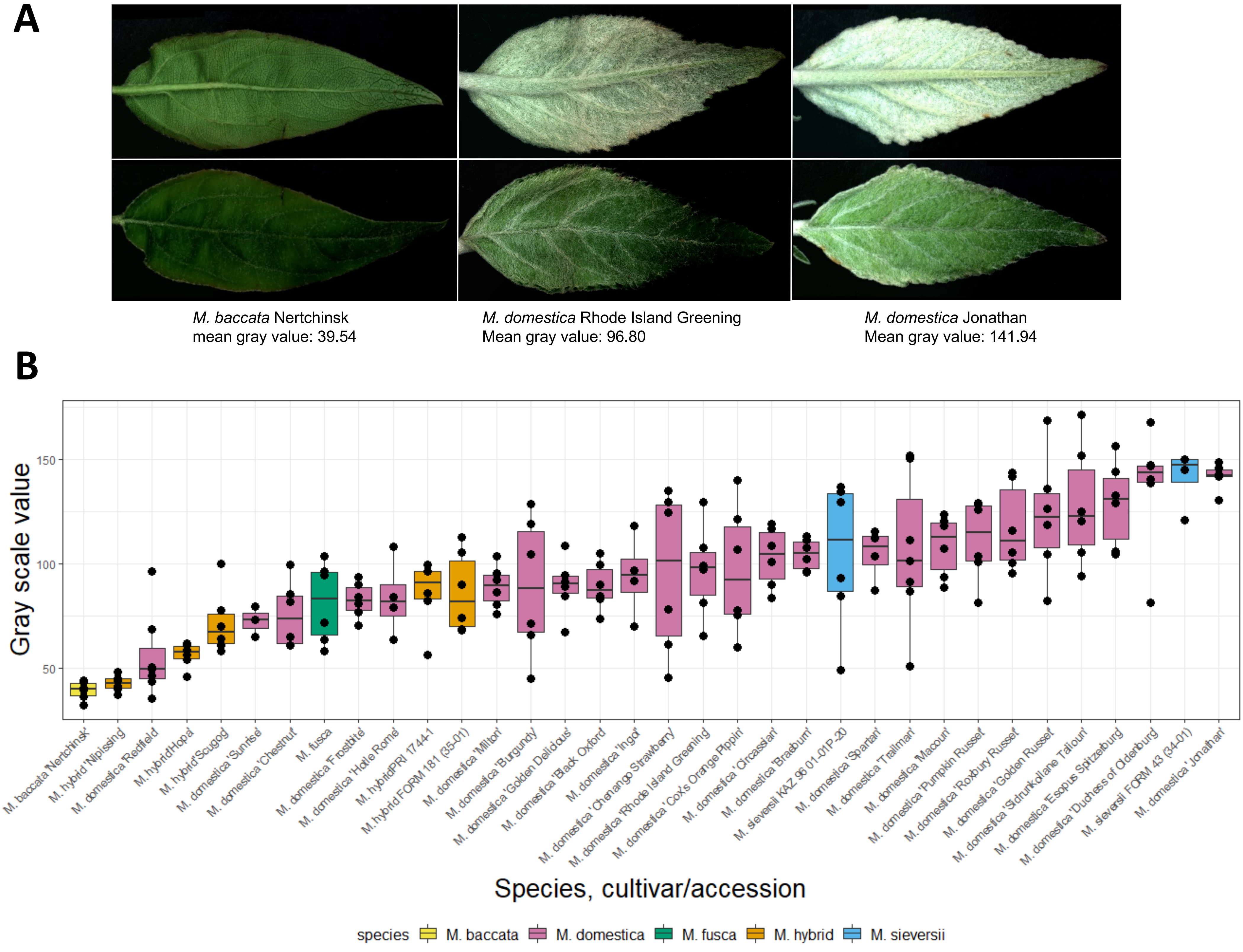

### Supplementary Fig. 5

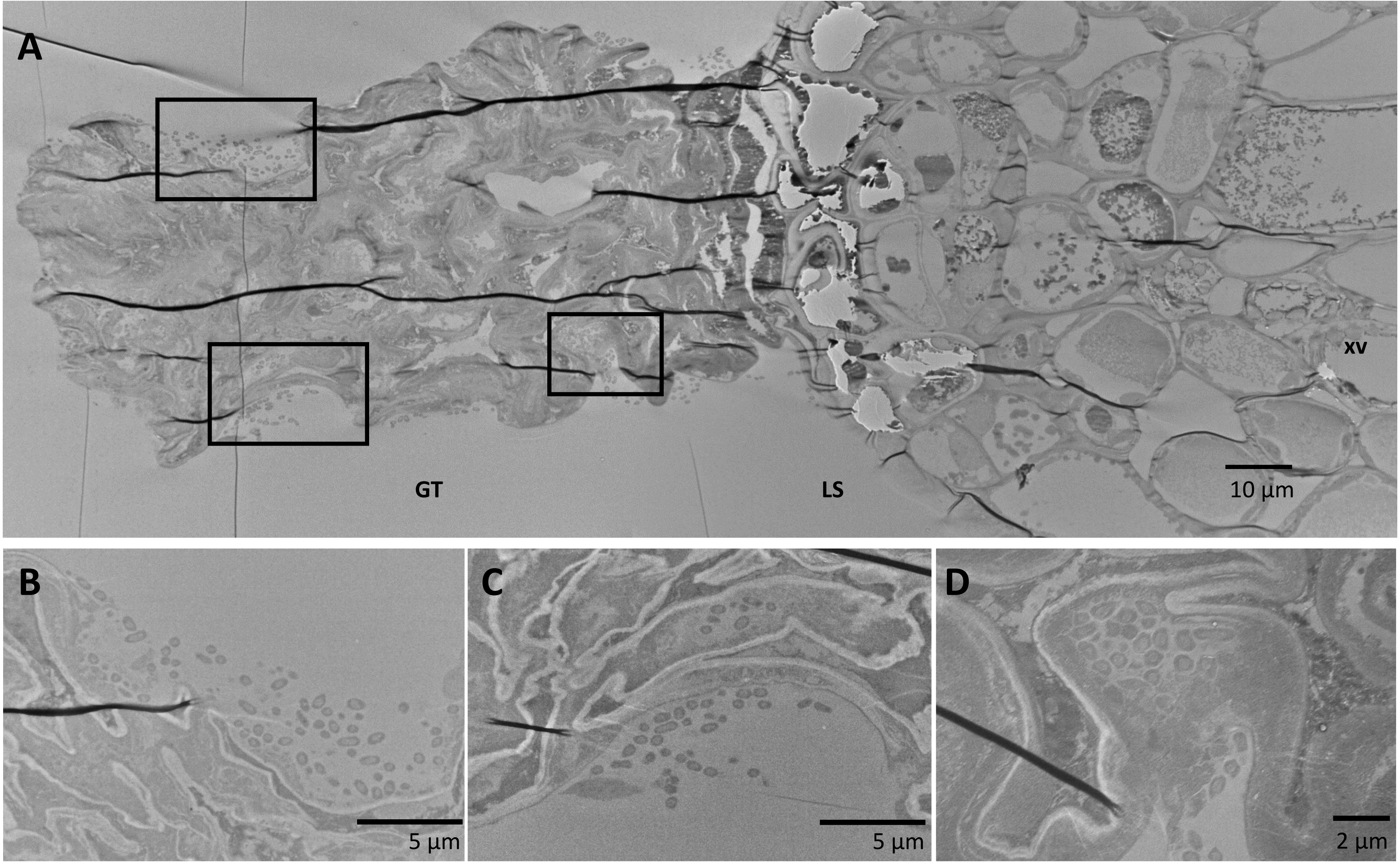
